# Support for the Adaptive Decoupling Hypothesis from Whole-Transcriptome Profiles of a Hypermetamorphic and Sexually Dimorphic Insect, *Neodiprion lecontei*

**DOI:** 10.1101/2019.12.20.882803

**Authors:** Danielle K. Herrig, Kim L. Vertacnik, Anna R. Kohrs, Catherine R. Linnen

**Affiliations:** Department of Biology, University of Kentucky, Lexington, KY, 40508, USA

**Keywords:** adaptation, antagonistic pleiotropy, metamorphosis, evolutionary constraint, transcriptomics, differential gene expression, holometabolous insects

## Abstract

Though seemingly bizarre, the dramatic post-embryonic transformation that occurs during metamorphosis is one of the most widespread and successful developmental strategies on the planet. The adaptive decoupling hypothesis (ADH) proposes that metamorphosis is an adaptation for optimizing expression of traits across life stages that experience opposing selection pressures. Similarly, sex-biased expression of traits is thought to evolve in response to sexually antagonistic selection. Both hypotheses predict that traits will be genetically decoupled among developmental stages and sexes, but direct comparisons between stage-specific and sex-specific decoupling are rare. Additionally, tests of the ADH have been hampered by a lack of suitable traits for among-stage comparisons and by uncertainties regarding how much decoupling is to be expected. To fill these voids, we characterize transcriptome-wide patterns of gene-expression decoupling in the hypermetamorphic and sexually dimorphic insect, *Neodiprion lecontei*. This species has three ecologically and morphologically distinct larval stages separated by molts, as well as a complete metamorphic transition that produces dimorphic adult males and females. Consistent with the ADH, we observe that: (1) the decoupling of gene expression becomes more pronounced as the ecological demands of developmental stages become more dissimilar and (2) gene-expression traits that mediate changing ecological interactions show stronger and more variable decoupling than expression traits that are likely to experience more uniform selection. We also find that gene-expression decoupling is more pronounced among major life stages than between the sexes. Overall, our results demonstrate that patterns of gene-expression decoupling can be predicted based on gene function and organismal ecology.

## Introduction

An estimated 80% of animal species have complex life cycles (CLCs) wherein metamorphosis separates two or more discrete, post-embryonic life stages (Wilbur, 1980). Although disagreements persist over diagnostic criteria, the general consensus is that metamorphosis involves an irreversible transformation in morphology that is typically accompanied by a pronounced change in ecology (Bishop et al., 2006). This change results in specialized stages optimized for distinct ecological niches (Benesh, Chubb, & Parker, 2013; Bishop et al., 2006; Ebenman, 1992; Istock, 1967; Moran, 1994). One explanation for the prevalence of CLCs is that independent adaptations at the different phases allow for optimal growth at some stages and optimal reproductive success at other stages (Bryant, 1969; Moran, 1994; Truman & Riddiford, 2019). Central to this explanation is the idea that pleiotropy creates genetic correlations across ontogeny that constrain evolution when traits beneficial for one stage are detrimental to another (Haldane, 1932).

The adaptive decoupling hypothesis (ADH) proposes that metamorphosis evolved as a mechanism for optimizing genetic correlations between life stages, thereby facilitating the independent evolution of traits when opposing selection pressures are experienced during different life stages (Ebenman, 1992; Haldane, 1932; Istock, 1967; Moran, 1994; Wigglesworth, 1954; Wilbur, 1980). A key prediction of the ADH is that traits that experience antagonistic selection across development will be genetically decoupled between distinct life stages and that more ecologically distinct life stages should have greater decoupling. To date, tests of this prediction have been mixed, with some studies supporting the ADH (Anderson, Scott, & Dukas, 2016; Blouin, 1992; Bonett & Blair, 2017; Goedert & Calsbeek, 2019; Hilbish, Winn, & Rawson, 1993; Jacobs et al., 2006; Loeschcke & Krebs, 1996; Medina, Vega-Trejo, Wallenius, Symonds, & Stuart-Fox, 2020; Parichy, 1998; Phillips, 1998; Saenko, Jerónimo, & Beldade, 2012; Sherratt, Vidal-García, Anstis, & Keogh, 2017; Wollenberg Valero et al., 2017), others refuting (Chippindale et al., 1998; Crean, Monro, & Marshall, 2011; Fellous & Lazzaro, 2011; Watkins, 2001; Wilson & Krause, 2012), and some with equivocal results (Aguirre, Blows, & Marshall, 2014; Helle, Johansson, Lederer, & Lind, 2010). However, many of these studies are limited by using only a few morphological traits and not taking stage-specific selection pressures into account when evaluating predictions of the ADH. Importantly, if metamorphosis is an adaptation for optimizing genetic independence, then the magnitude of trait decoupling should depend on the strength of antagonistic selection. Testing this prediction of the ADH will require quantifying decoupling for large and diverse collections of traits that vary in the extent to which they experience antagonistic selection.

Whole-transcriptome gene-expression data obtained from multiple life stages provide an ideal collection of traits for evaluating the extent to which patterns of genetic decoupling fit predictions of the ADH (Collet & Fellous, 2019). First, because all stages of the life cycle must be encoded by a single genome, dramatic phenotypic changes that accompany metamorphosis must be mediated by changes in gene expression. Second, transcriptomes provide a large number of quantitative traits, all measured in comparable units of gene expression, that can be readily compared across life stages. Third, the genes included in a transcriptome cover a wide range of biological functions that should vary somewhat predictably in the extent to which they experience antagonistic selection across the life cycle. Following the logic of the ADH, this variation in selection should generate predictable variation in gene-expression decoupling. For example, genes involved in basic cellular functions (i.e., housekeeping genes) should be more genetically coupled than genes that mediate ecological interactions that change across the life cycle.

Transcriptome-wide patterns of decoupling should also vary with the magnitude of the ecological change accompanying metamorphosis. The most extreme metamorphic transformations occur in holometabolous insects, whose defining characteristic is a distinct pupal stage that separates two completely different body plans (Gilbert, Tata, & Atkinson, 1996; Heming, 2003; Kristensen, 1999). Importantly, this profound transformation enables one stage to be optimized for feeding and growth (the larval stage) and a second stage for dispersal and reproduction (the adult stage). The genetic independence of larval and adult traits proposed by the ADH may explain, in part, why holometabolous insects are one of the most evolutionarily successful and diverse lineages on the planet (Ebenman, 1992; Haldane, 1932; Istock, 1967; Moran, 1994; Rainford, Hofreiter, Nicholson, & Mayhew, 2014; Truman & Riddiford, 2019; Wigglesworth, 1954; Wilbur, 1980). In some holometabolous lineages, pronounced ecological and morphological transformations also occur between successive larval instars. This phenomenon, which has been dubbed hypermetamorphosis (Belles, 2011), provides a valuable opportunity to test the prediction that transcriptome-wide levels of genetic decoupling between life stages will increase with the dissimilarity of the fitness landscapes to which they are adapting.

Looking beyond metamorphosis, the rationale underlying the ADH applies more generally to any scenario in which a single genome expresses multiple distinct phenotypes that are subject to opposing selection pressures (Collet & Fellous, 2019). Arguably the best-studied scenario of genetic decoupling evolving in response to antagonistic pleiotropy involves alternative phenotypes of a single life stage: adult males and adult females. Just as stage-limited gene expression can reduce genetic correlations across life stages of organisms with CLCs, sex-biased gene expression can enable the independent evolution of male and female traits, leading to the evolution of sexual dimorphism (Assis, Zhou, & Bachtrog, 2012; Ellegren & Parsch, 2007; Parsch & Ellegren, 2013; Perry, Harrison, & Mank, 2014; Proschel, Zhang, & Parsch, 2006). An important distinction between CLCs and sexual dimorphism is that only in CLCs must all alternative phenotypes (i.e., life stages) have non-zero fitness for a novel pleiotropic allele to spread in a population (Collet & Fellous, 2019). More generally, for the same net fitness difference between alternative phenotypes, selection against an allele with opposing fitness effects in different life stages may be stronger than selection against an allele with opposing fitness effects in different sexes. For this reason, trait decoupling should be more pronounced for ecologically distinct life stages than for different sexes.

To date, only a handful of studies have evaluated the prediction that gene-expression traits will be decoupled across metamorphic boundaries as it pertains to the adaptive decoupling hypothesis (Fellous & Lazzaro, 2011; Jacobs et al., 2006; Saenko et al., 2012; Wollenberg Valero et al., 2017). Furthermore, to our knowledge, no study has evaluated whether gene-expression decoupling varies predictably with the type of life-cycle transition or gene function, and few studies have directly compared patterns of sex-biased and stage-biased gene expression (but see (Ometto, Shoemaker, Ross, & Keller, 2011; Perry et al., 2014)). To these ends, we take advantage of a hypermetamorphic and sexually dimorphic species of insect with a well-characterized ecology and annotated genome, the redheaded pine sawfly *(Neodiprion lecontei,* order: Hymenoptera; family: Diprionidae) (Figure 1).

**Figure 1:**
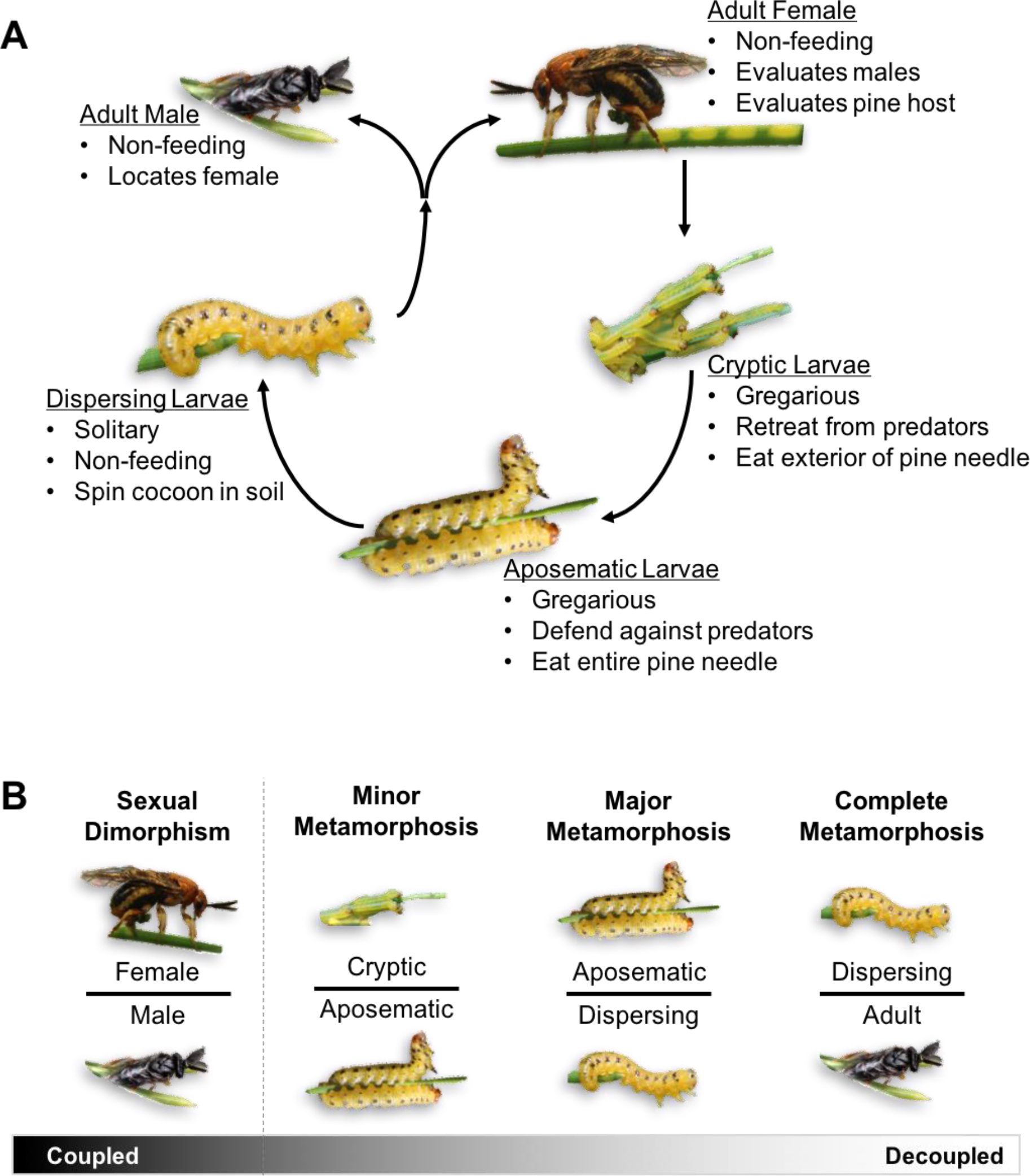
Ecological changes during *N. lecontei* development generate predictions regarding geneexpression decoupling between the sexes and across different metamorphic transitions. (A) The life cycle of the red-headed pine sawfly includes two distinct feeding larval stages, a non-feeding larval stage, and sexually dimorphic adults. Bullet points highlight key behaviors that change across development. (B) Based on this life cycle, metamorphic transitions include: minor metamorphosis (between the two feeding larval stages), major metamorphosis (between the feeding and non-feeding larval stages), and complete metamorphosis (between larval stage and adult). The gradient represents our prediction that gene expression will be most coupled between the feeding larval stages and the adult sexes while it is most decoupled for stages separated by complete metamorphosis.

In addition to the complete metamorphic event that occurs during the pupal stage, there are two metamorphic transitions that occur within the larval stage of the redheaded pine sawfly that result in pronounced changes in coloration and behavior (Atwood & Peck, 1943; Coppel & Benjamin, 1965; Linnen, O’Quin, Shackleford, Sears, & Lindstedt, 2018) and references therein) (Figure 1A). The first metamorphic transition is a shift from a “cryptic” to an “aposematic” feeding larval morph and is less dramatic than the other transitions (hereafter, “minor metamorphosis”). The cryptic morph is lightly pigmented, ingests only the exterior of pine needles while avoiding the toxic resinous core, and retreats to the base of the needle when predators are near. By contrast, the aposematic morph is heavily pigmented, ingests the entire needle, and sequesters the toxic pine resins for use in dramatic anti-predator defensive displays. A more striking transition occurs when the aposematic morph molts into a “dispersing” final instar (hereafter, “major metamorphosis”). The dispersing morph is solitary, non-feeding, less intensely pigmented, and migrates to the litter or soil to spin a cocoon. Complete metamorphosis occurs within the cocoon. The non-feeding adult stage is dedicated entirely to reproduction. Sexually dimorphic adults are highly specialized for sex-specific tasks. Males are excellent fliers and use bipectinate antennae to detect female pheromones from considerable distances. In contrast, females remain near the cocoon eclosion site and use serrate antennae to search for suitable oviposition sites in *Pinus* needles (Anderbrant, 1993). Like most hymenopterans, *N. lecontei* females lay a combination of fertilized and unfertilized eggs that will develop into diploid females and haploid males, respectively.

The hypermetamorphic life cycle of the redheaded pine sawfly, the wealth of natural history data for this species, and the logic of the ADH enables us to make *a priori* predictions about how levels of genetic decoupling (inferred here from the magnitude of differential gene expression) will vary among genes categories, developmental stages, and sexes. We predict that: 1) The extent of gene-expression decoupling should increase with the ecological dissimilarity of the life-stages (Figure 1B). 2) Across the transcriptome, the most pronounced gene-expression decoupling will be observed for genes that mediate ecological changes across development. 3) Because traits expressed in different individuals (sexes) may experience weaker selection for decoupling than traits expressed in multiple life stages of a single individual, we predict trait decoupling between the sexes will be less extreme than that observed between metamorphic events (Figure 1B). To test these predictions, we generated expression data for 9,304 genes via whole-transcriptome sequencing of males of each *N. lecontei* life stage and adults of both sexes.

## Materials and Methods

### Tissue dissection in lab-reared *N. lecontei*

To obtain a large number of individuals for tissue dissection, we established laboratory colonies of *N. lecontei* from a single population of larvae collected in 2016 from a *Pinus mugo* bush in Lexington, KY, USA (38.043602°N, 84.496549°W). We reared colonies on *Pinus* foliage using standard rearing protocols (Bendall, Vertacnik, & Linnen, 2017; Harper, Bagley, Thompson, & Linnen, 2016) for a minimum of one full generation before collecting tissues. To ensure that all larvae were haploid males, we collected male tissues (all larval tissues and adult males) from the progeny of virgin females. We collected adult female tissues from the progeny of mated females. To obtain larval tissues, we collected different instars of haploid male larvae from rearing boxes containing *Pinus* foliage. We immediately placed live larvae on a petri dish containing frozen 1X PBS buffer, removed the heads from the bodies, and flash-froze both tissues on dry ice prior to storage at −80°C. To obtain adult tissues, we stored adults at 4°C (a temperature that extends lifespan) for up to 20 days until tissues could be collected. Before harvesting tissues, we warmed adults to room temperature and enclosed them in a mesh cage containing three *Pinus* seedlings for a minimum of 15 minutes to induce normal host- and mate-seeking behavior. Then, to sedate adults prior to dissection, we enclosed each individual in a deli cup and exposed it to dry ice for approximately 1 minute. We placed the sedated adult on frozen 1X PBS buffer to harvest six tissues: antennae, mouthparts, heads (sans antennae and mouthparts), legs, thorax, and the terminal segment of the abdomen (which includes the ovipositor in females or copulatory organ in males). We flash-froze these tissues and stored them at −80°C until RNA extraction.

### RNA extraction, library preparation, and sequencing

We extracted RNA from all tissues with RNeasy Plus Micro and Mini Kits (Qiagen, Germantown, Mayland). To account for family-level variation in gene-expression traits, each RNA extraction contained tissues from a single lab-reared family (each represented by 2-18 individuals, depending on the tissue). Prior to starting standard kit protocols, we disrupted tissues using a TissueLyser LT bead mill (Qiagen, Germantown, Mayland) and 5mm stainless steel beads. We disrupted tissues for up to four 30-sec periods of 60 oscillations/sec, placing tissues on dry ice between periods to ensure that the RNA did not degrade during the disruption process. We then quantified RNA using Quant-iT RNA Assay Kits (Invitrogen by ThermoScientific, Waltham, Massachusetts) and assessed quality using a 2100 Bioanalyzer and RNA 6000 Nano Kits (Agilent, Santa Clara, California).

To prepare RNA-seq libraries for Illumina sequencing, we used a TruSeq Stranded mRNA High Throughput Kit (Illumina, San Diego, California), following manufacturer protocols. For larval tissues, we made each library from RNA extracted from members of a single family. For adult tissues, which required more individuals to obtain sufficient material, we used 1-3 families per library (with equal RNA contributions from each family). For most tissues, we made libraries for four biological replicates, with different families in each replicate. The exceptions were male copulatory organs (for which only 3 high quality biological replicates could be produced) and male thoraxes (for which only 2 high quality biological replicates could be produced), resulting in a total of 77 libraries (Supplemental Table 1). We quantified resulting cDNA using Quant-iT DNA Assay Kits (Invitrogen by ThermoScientific, Waltham, Massachusetts) and evaluated quality with a 2100 Bioanalyzer and DNA 1000 Kits (Agilent, Santa Clara, California). We then pooled libraries that passed quality inspection and sequenced this pool with 150-bp paired-end reads on an Illumina HiSeq4000 at the University of Illinois at Urbana-Champaign Roy J. Carver Biotechnology Center.

### Processing of RNAseq data to produce a provisionally annotated *de novo* transcriptome

To de-multiplex and quality-trim raw reads, we used trimmomatic (Bolger, Lohse, & Usadel, 2014) and a cut-off score of 30. We also removed any remaining TruSeq indexed universal adapter present at the beginning of reads using fastx_clipper (http://hannonlab.cshl.edu/fastx_toolkit/). After filtering, we used tophat2 (Kim et al., 2013) to map retained reads to the *N. lecontei* version 1.0 assembly (Vertacnik, Geib, & Linnen, 2016). We then used a these mapped reads to produce a genome-guided *de novo* transcriptome assembly using Trinity (Haas et al., 2013). Because TRINITY is known to overestimate the actual number of contigs present in transcriptomes (https://github.com/trinityrnaseq/trinityrnaseq/wiki), we performed additional filtering to retain meaningful contigs. First, we used CD-HIT (W. Li & Godzik, 2006) to create non-redundant contigs from the TRINITY contigs (minimum of 200bp). Next, we used Bowtie2 (Langmead & Salzberg, 2012) to map the reads to the non-redundant contigs and RSEM (B. Li & Dewey, 2011) to identify contigs with at least one transcript per million in at least two biological replicates.

To functionally annotate contigs that were retained after filtering, we performed two sets of BLASTx (Altschul, Gish, Miller, Myers, & Lipman, 1990) searches. First, we BLASTed these transcripts against the predicted *N. lecontei* non-redundant proteins available on NCBI and a set of manually curated *N. lecontei* OBPs, ORs, and GRs (Vertacnik *et al*. in prep). For transcripts that did not map with 90% identity to putative *N. lecontei* genes, we performed an additional BLAST search against the insect non-redundant protein database (Pruitt, 2004) (e-value of 0.001). In all cases, we selected the best BLAST match as a provisional annotation for these transcripts.

### Testing predictions of the ADH

If gene-expression traits are “adaptively decoupled” there should be variable and predictable patterns of decoupling across the transcriptome. At a phenotypic level, gene-expression traits that are highly decoupled across development are expected to exhibit: (1) pronounced differences in expression levels, (2) minimal quantitative genetic correlations, and (3) independent evolutionary trajectories across phylogenies (Collet & Fellous, 2019; Medina et al., 2020; Wollenberg Valero et al., 2017). At a genetic level, variation in adaptively decoupled gene-expression traits within and between species should be attributable to mutations with minimal pleiotropy across life stages. Ideally, each of these four patterns (differential expression, reduced quantitative genetic correlations, macroevolutionary independence, and reduced levels of pleiotropy) would be evaluated across the entire transcriptome. However, even with current sequencing technologies and novel tools for functional genomics, transcriptome-wide analyses would be prohibitively expensive for the large sample sizes needed for quantitative genetic, comparative phylogenetic, and forward or reverse genetic approaches. Therefore, as a first step to quantifying transcriptome-wide patterns of gene-expression decoupling across multiple metamorphic transitions, we used differential expression between life stages and sexes of a single species as our measure of decoupling.

To compare levels of differential expression across different metamorphic transitions and among different types of genes, we first used bowtie2 (Langmead & Salzberg, 2012) to align reads to the *de novo N. lecontei* transcriptome described above. We then estimated the abundance of each transcript using RSEM via the Trinity package utility program align_and_estimate_abundance.pl (Haas et al., 2013; B. Li & Dewey, 2011). Using the utility program abundance_estimates_to_matrix.pl (Haas et al., 2013) we created a complete count matrix of transcript abundance. This program also produces matrices of normalized gene expression for comparisons of relative abundances (transcripts per million) as well as correcting for highly expressed genes to obtain absolute abundances (TMM, third-quartile normalization) (Haas et al., 2013).

To determine whether gene-expression decoupling changes predictably across different types of metamorphic transitions, we compared whole-transcriptome profiles of different life stages and sexes in two ways. First, to visualize overall similarity or dissimilarity of gene expression across all tissues and life stages, we conducted a principle component analysis (PCA) using the PtR scripts within the Trinity package (Haas et al., 2013). Second, to quantify how decoupling changes across different metamorphic boundaries and between the sexes, we compared the number of differentially expressed genes (DEGs). To determine the number of DEGs, we used the Trinity utility program run_DE_analysis.pl to implement DESeq2 (Haas et al., 2013; Love, Huber, & Anders, 2014) and obtain statistics of differential expression between different tissue types and different life stages. To account for multiple testing this program utilized the Benjamini-Hochberg correction, requiring an adjusted p-value *(padj)* of 0.05 or less to be considered significantly differentially expressed. We compared the numbers of differentially expressed genes between metamorphic transitions using Fisher’s exact test and accounted for multiple testing using “RVAideMemorire” package (v. 0.9-70, implemented in R 3.5.0).

To determine whether genes that mediate changing ecological interactions show stronger and more variable levels of gene-expression decoupling, we used both ad hoc and a priori approaches to evaluate candidate genes. For our ad hoc approach, we compiled lists of the top differentially expressed genes (i.e., lowest p-values) between each of the three metamorphic transitions and between the sexes using the output of DESeq2. For each comparison, we identified the top 10 genes that were upregulated in each tissue and in each stage/sex and asked whether these lists contained any genes that are likely to mediate ecological interactions. Based on the ecology of *N. lecontei* (Figure 1), we were specifically looking for genes involved in digestion, detoxification, pigmentation, gregarious behavior, chemosensation, immune function, and reproduction. We first asked whether any of our top differentially expressed genes corresponded to candidate genes in existing manually curated gene datasets for *N. lecontei* (chemosensation, detoxification, and immunity genes: (Vertacnik et al., in prep); pigmentation genes: (Linnen et al., 2018)). For remaining genes, we identified the closest *Drosophila* orthologue and used the gene ontology (GO) term database to determine the likely function for each gene. When no *Drosophila* orthologue could be identified or the orthologue did not have a predicted GO function, we used UniProt to predict the possible function of the gene.

For our a priori approach, we compared patterns of gene-expression decoupling for two comparably sized sets of genes that we expected to be under drastically different selection regimes. The first was set of manually curated chemosensory genes (including odorant receptors, gustatory receptors, odorant-binding proteins; (Vertacnik et al., in prep).), which are expected to experience more antagonistic selection and, therefore, exhibit more extreme and more variable decoupling. The second category consisted of a similarly sized family of housekeeping genes (the *ribosomal protein L* genes, hereafter RPLs), which are expected to show coupled expression across sexes and life stages. To visualize how the degree of gene-expression decoupling of our “high” and “low” decoupling categories compare to the rest of the transcriptome, we first condensed each gene in the transcriptome to a single expression value per stage/sex. For each gene, we calculated the log2 of the average normalized expression level across all tissues and replicates for each life stage. We then overlaid RPL and chemosensory gene-expression values on transcriptome-wide values for each metamorphic event and between the sexually dimorphic adults. To determine whether the distribution of expression differences between the stages differs between RPLs and chemosensory genes, we first calculated the absolute value of the average log2-fold change for each comparison with DESeq2. Then, we used non-parametric Mann-Whitney U tests to compare the two groups of genes within each sexual/metamorphic comparison.

Because our analyses revealed highly variable patterns of decoupling across chemosensory genes (see below), we went a step further to identify chemosensory genes with the highest and lowest levels of decoupling. We investigated variation in decoupling among chemosensory genes in two ways. First, we produced heatmaps to visualize variation in patterns of decoupling across stages/sexes across individual chemosensory genes. Because there was drastic variation in the maximum expression of chemosensory genes, we first used the scale function so that the expression of so that the total expression of each gene was equal and the heatmap function to visualize the normalized expression levels using R 3.3.2 (Team, 2016). Finally, we used custom Python (v3.5.1) scripts to identify chemosensory genes that had expression levels in the top 10th percentile in the feeding larvae, adult males, and adult females to further pinpoint genes with coupled or decoupled expression across life stages.

## Results

### Sequencing and transcriptome assembly

In total, we obtained over 787 million high-quality paired-end 150-bp reads, with an average of 39.4 million reads per tissue type (Supplemental Table 1; accession numbers will become available upon publication). After filtering, we retained over 766 million reads (Supplemental Table 2). An average of 88.4% of these reads mapped to the *N. lecontei* genome. Our genome-guided *de novo* transcriptome assembly identified 132,243 contigs. After further filtering, we retained 58,353 contigs. Across our 77 libraries, average mapping rate to these contigs was 85.5% (range: 74.9-91.6%). The provisionally annotated transcriptome contained 16,714 contigs representing 9,304 predicted insect genes. Using these 16,714 contigs as the transcriptome, a high mapping rate was maintained across all libraries (mean: 76.7%; range: 60.5-88.6%).

### Decoupling of gene-expression profiles increases with ecological dissimilarity of *N. lecontei* life stages

Consistent with our prediction that differences in gene-expression traits would be most pronounced between ecologically dissimilar life stages (Figure 1B), stages separated by major and complete metamorphosis were clearly distinct as shown in the first principle component (PC1) of our principle component analysis (Figure 2A). By contrast, there were no clear distinctions between cryptic and aposematic larvae or between males and females along either of the first two gene-expression PCs. PC2 primarily separated tissues within each life stage (Figure 2A). Along this axis, gene expression in larval and adult heads was clearly distinct from expression in other tissues, as was expression in the adult antennal tissues. Based on these results, we analyzed gene-expression decoupling among stages/sexes in two ways: all tissues combined and heads only (Supplemental Tables 3 and 4, respectively).

**Figure 2:**
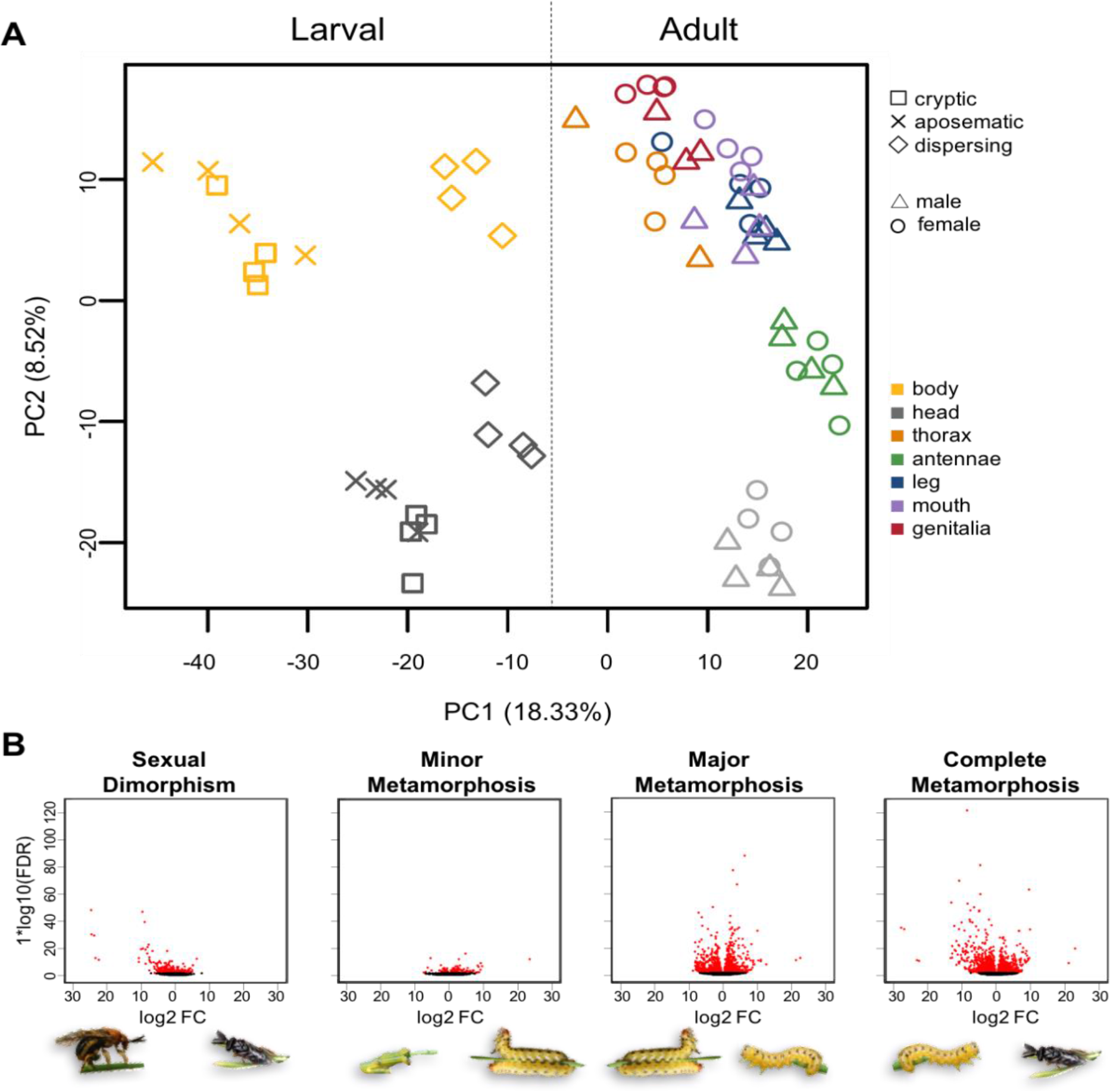
Transcriptome-wide patterns of gene-expression decoupling are consistent with predictions of the ADH. (A) Principle component analyses of 9,304 putatively annotated genes for the head (gray) and body (yellow) tissues of cryptic (✉), aposematic (×), and dispersing (O) larvae as well as the head (grey), antenna (green), mouthpart (purple), leg (navy), genitalia (red), and thoracic (orange), tissues of adult males (Δ) and females (o). (B) Volcano plots with the log2-fold change of gene expression and log10 FDR-adjusted P-values between the sexually dimorphic adults and between each of the metamorphic transitions, arranged according to predictions as in Figure 1B. Each point represents a single gene; red points are significantly differentially expressed between the stages/sexes.

Regardless of whether we looked at all tissues or heads only, both the magnitude of differential expression (log fold-change) and significance (Benjamini-Hochberg adjusted p-value) increased in accordance with ADH predictions across metamorphic transitions (Figure 2B for all tissues). Specifically, the percentage of DEGs increased with ecological dissimilarity of the life stages compared: 2.2% of genes were differentially expressed between cryptic and aposematic larvae, 24.7% between aposematic and dispersing larvae, and 25.1% between dispersing larvae and adult males (Supplemental Table 3; heads only: 2.9%, 17.6%, and 31.1%, respectively, Supplemental Table 4). The proportion of DEGs increased concurrently with the extremity of metamorphosis (Fisher’s exact tests; *P* < 1 x 10-20) except for the all-tissue dispersing larvae vs. male comparison (*P =* 0.59).

Despite males and females being more morphologically dissimilar than the feeding and dispersing larval stages, the percentage of DEGs between sexually dimorphic adults (all tissues: 7.0%, Supplemental Table 3; head only: 6.5%, Supplemental Table 4) was far less than that observed across both the major and complete metamorphic transitions (Fisher’s exact tests; *P* < 1 x 10-208)(Figure 2B). In contrast, we observed a higher percentage of DEGs between the sexes than between the two feeding larval stages (Fisher’s exact tests; *P* < 1.0 x 10-20). Taken together with the PCA, these results indicate that gene-expression decoupling between the sexes is more or less on par with the minor metamorphic transition between cryptic and aposematic larvae and far less pronounced than the decoupling observed between more extreme life-stage transitions. Overall, these transcriptome-wide analyses support the ADH prediction that genetic decoupling increases as the ecological demands of the different life stages become more dissimilar, as well as the prediction that genetic decoupling tends to be more pronounced between developmental stages than between the sexes.

### Variation in decoupling among different types of gene-expression traits reflects changes in ecology

Consistent with our prediction that genes that mediate changing ecological interactions will exhibit the most pronounced decoupling, many of the top differentially expressed genes between life stages/sexes corresponded to candidate genes that were related to their ecological differences (Supplemental Table 5). Among the top differentially expressed genes between cryptic and aposematic larvae were genes potentially involved in immunity, metabolism and detoxification. For example, *esterase FE4-like* (elevated in bodies of aposematic feeding larvae), is a gene thought to be involved in resistance to organophosphate insecticides and may be essential to ingesting and sequestering toxic pine resins. The *esterase FE4-like* gene was also among the top genes downregulated in the non-feeding dispersing larvae. The change from an aposematic feeding larva to a dispersing larva was also accompanied by a pronounced decrease in expression of *Cameo2,* a gene that is thought to play a role in carotenoid-based pigmentation (Y. Li et al., 2014) and has been linked to larval body color in *N. lecontei* via QTL mapping (Linnen et al., 2018). Putative pigmentation and detoxification genes were also among the top differentially expressed genes between larvae and adult males (Supplemental Table 5).

The top differentially expressed genes between males and females for each tissue appear to be related to the sex-specific tasks of the adults (Supplemental Table 5). For example, females, which must properly provision their eggs, had elevated expression for a *vitellogenin-like* gene, which is associated with egg yolk formation, hormone regulation, lifespan, and foraging behavior (Ihle, Page, Frederick, Fondrk, & Amdam, 2010; Munch, Ihle, Salmela, & Amdam, 2015; Nunes, Ihle, Mutti, Simoes, & Amdam, 2013; Seehuus, Norberg, Gimsa, Krekling, & Amdam, 2006; Wheeler, Ament, Rodriguez-Zas, & Robinson, 2013) There was also increased expression for many *major royal jelly* protein family genes, known to play a role reproductive maturation and tending to young (Buttstedt, Moritz, & Erler, 2013; Dobritzsch, Aumer, Fuszard, Erler, & Buttstedt, 2019; Drapeau, Albert, Kucharski, Prusko, & Maleszka, 2006). Among the genes with unusually high expression in males were three chemosensory genes *(OR54, OR22,* and *OBP9)* that were particularly high in the antennae and may play a role in mate finding.

### Chemosensory genes vary in decoupling across life stages

Consistent with our a priori predictions, the magnitude of decoupling for manually curated chemosensory genes was highly variable (Figure 3A). While several chemosensory genes had similar levels of expression across life stages (e.g. those falling along the dotted lines), others were among the most decoupled genes in the transcriptome (e.g., those falling at the edges of the transcriptome-wide cloud of points). This contrasted with a family of housekeeping genes that have a similar family size—the *ribosomal protein L* genes (RPLs). As expected, these genes were highly coupled across development and had nearly identical expression levels in comparisons between the all sexes or stages (e.g., those that fall on the dotted line). Figure 3B directly compares variation in decoupling for these two categories of genes. Consistent with the ADH, gene-expression decoupling for chemosensory genes was both more variable (i.e., wider distribution) and significantly higher than RPLs in 3 out of 4 comparisons (Figure 3B and Supplemental Table 4; Mann-Whitney U test *P* < 1.0 x 10-3). The exception was the aposematic vs. dispersing larvae comparison representing the major metamorphic transition (*P* = 0.90). Overall, these results are consistent with a scenario in which selection has favored coupling for some chemosensory genes and stage-specific decoupling for others.

**Figure 3:**
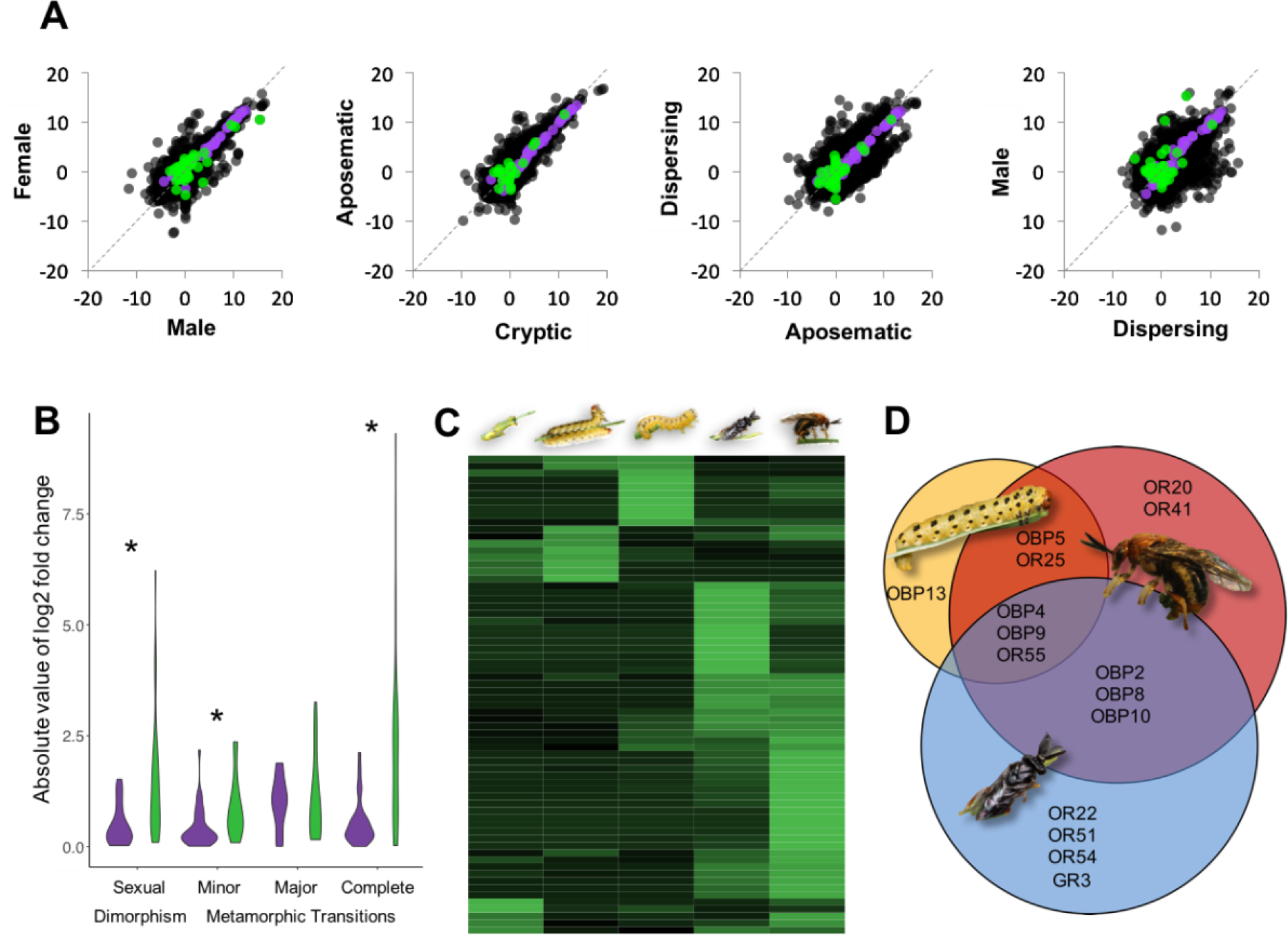
Compared to a group of house-keeping genes, chemosensory genes have higher and more variable levels of expression decoupling across development and between the sexes. (A) Correlation plots of the sexually dimorphic adults and each metamorphic stages for all genes in the transcriptome (black points), Ribosomal protein L genes (purple points), and chemosensory genes (green points). Each point represents log2 of the average expression level across all tissues/replicates for each life stage/sex. The dotted line represents equal expression in the stages/sexes being compared; deviation from this line represents gene-expression decoupling. (B) Violin plots showing the distributions of the absolute value of the log2-fold change between each comparison (calculated as in Figure 2B) for Ribosomal protein L genes (purple) and chemosensory genes (green). Comparisons are for adult male vs. adult female (Sexual Dimorphism) and the three metamorphic transitions: cryptic vs. aposematic larvae (Minor), aposematic vs. dispersing larvae (Major), and dispersing larva vs. adult male (Complete). Asterisks indicate comparisons for which decoupling is significantly higher for chemosensory genes than for RPL genes. (C) Heatmap of expression for each chemosensory gene in 4 male life stages and in adult females (from left to right: cryptic larvae, aposematic larvae, dispersing larvae, adult male, and adult female). Expression is adjusted such that each row has the same total amount of expression to reveal expression patterns for genes expressed at relatively low levels. Whereas some genes have high expression in only one life stage, others are more widely expressed. (D) Venn diagram of chemosensory genes with expression levels in the top 10% of all genes expressed in each tissue for feeding larvae, adult males, and adult females. Genes include all olfactory receptors (ORs), gustatory receptors (GRs), and odorant binding proteins (OBPs). Whereas some chemosensory genes exhibit highly sex- and stage-specific expression, others are less decoupled and expressed in multiple stages/sexes.

To gain additional insight into how chemosensory gene expression changes across the life cycle, we created a heat map of stage-specific expression for each gene (Figure 3C). Whereas some chemosensory genes were expressed only during a single life stage, others were expressed at moderate to high levels in multiple life stages. We further analyzed these genes by identifying those that were in the highest 10% of expression for each stage/sex. This analysis revealed several chemosensory genes that were highly expressed in a single stage/sex while others were highly expressed in multiple stages and/or both sexes (Figure 3D and Supplemental Table 6). Together, these patterns provide clues into chemosensory gene function. For example, chemosensory genes that were highly expressed by both feeding larvae and females (*OBP5* and *OR25*) could be involved in detecting host-plant cues. Likewise, genes highly expressed in both sexes of reproductive adults *(OBP2, OBP8,* and *OBP10*) may play roles in conspecific mate recognition. Genes expressed by all three stages (*OBP4*, *OBP9*, and *OR55*) may be important for detecting conspecific or host cues that are critically important throughout the life span. In contrast, genes with high expression in only a single stage may only be essential for a single task or ecological pressure. For example, *OBP13* is expressed at high levels in feeding larvae with low levels of expression in dispersing larvae an almost no expression in adults. Each adult sex also has specific chemosensory genes with *OR20* and *OR41* expressed almost exclusively in female antennae while *OR22*, *OR51*, and *OR54* expressed almost exclusively in male antennae. Notably, although all larvae were males, we did not identify any genes that were highly expressed in larval and adult males to the exclusion of adult females, suggesting once again that ecology is a better predictor of gene-expression patterns than sex in this species.

## Discussion

According to the ADH, complex life cycles are pervasive in nature because they facilitate the independent evolution of traits expressed at different life stages, thereby removing constraints that may otherwise slow or prevent adaptation to novel environments (Moran, 1994). Evidence for a key prediction of the ADH—that traits are decoupled across metamorphosis—has been mixed ((Aguirre et al., 2014; Collet & Fellous, 2019; Fellous & Lazzaro, 2011; Freda, Alex, Morgan, & Ragland, 2017; Saenko et al., 2012; Sherratt et al., 2017) and references therein). Here we argue that if metamorphosis is an adaptation for optimizing genetic correlations across life stages, genetic decoupling may evolve only for traits that experience antagonistic selection across ontogeny. In other words, the ADH does not necessarily predict that all traits will be decoupled, but rather that trait decoupling should be somewhat predictable given sufficient knowledge of stage-specific selection pressures. Therefore knowledge of how selection pressures may differ on traits across development is essential to predicting and interpreting patterns of decoupling for individual gene-expression traits with respect to testing the ADH.

To evaluate the predictability of trait decoupling in light of the ADH, we took advantage of a well-characterized hypermetamorphic insect, the redheaded pine sawfly (Figure 1) to generate *a priori* predictions regarding the decoupling of gene-expression traits. Consistent with our predictions, we found that: 1) ecological differences between life stages enabled us to accurately predict how transcriptome-wide levels of gene-expression decoupling vary across development (Figure 2). 2) Levels of decoupling vary among functionally different genes in the transcriptome, with ecologically relevant genes being some of the most highly and variably decoupled genes between life stages (Supplemental Tables 3-5 and Figure 3). 3) Decoupling of gene-expression traits tended to be more pronounced between developmental stages than between sexes (Figures 2 and 3). Here, we compare our results to other studies of trait decoupling under ontogenetically and sexually antagonistic selection and, in light of this body of work and limitations of our own study, make suggestions for future research on the ADH.

### The predictably variable decoupling of some ecologically important traits

If metamorphosis facilitates optimization of genetic correlations, then patterns of trait decoupling should depend on how changing ecological selection pressures align with genetic correlations across development (Collet et al. 2019). Variation in selection pressures experienced by different traits should generate corresponding variation in decoupling patterns. If selection optimizes gene-expression traits, the extent of gene-expression decoupling across life stages/sexes should vary even among genes belonging to the same narrow functional category depending on precise ecological roles (e.g., (Fellous & Lazzaro, 2011)). Consistent with this expectation, we observed a wide range of decoupling patterns for expression levels of different chemosensory genes (Figure 3). In some cases, antagonistic selection may have favored decoupling of gene expression traits—for example, perhaps males and females express different chemosensory genes to optimize mate- and host-finding, respectively. In other cases, selection may have favored positive genetic correlations between host-use traits expressed at different life stages (Agrawal & Stinchcombe, 2009; Vertacnik & Linnen, 2017). For example, in sawflies, each life stage interacts with the host plant and these stages may express some chemosensory genes across development (coupled expression) to monitor important host chemical cues (Figure 1A, Figure 3D). Beyond sawflies, numerous studies of stage-specific gene expression in many other insect taxa have found trends very similar to those reported here, with chemosensory genes exhibiting both high and low levels of gene-expression decoupling (Colgan et al., 2011; Etges et al., 2015; Harker et al., 2013; Koutsos et al., 2007; Lee et al., 2016; Sayadi, Immonen, Bayram, & Arnqvist, 2016; Schönbach et al., 2013; H. Yang et al., 2018; Zhang, He, & Wang, 2016). Thus, high variance among chemosensory genes in the degree of coupling/decoupling across development may be a common outcome of adaptive gene expression evolution (Vertacnik & Linnen, 2017).

Mixed decoupling patterns have also been observed for other types of ecologically important gene-expression traits. One such example comes from a 2011 study that investigated expression decoupling of two antimicrobial peptide (AMP) genes between *Drosophila melanogaster* larvae and adults (Fellous & Lazzaro, 2011). Using a quantitative genetic approach, Fellous and Lazarro (2011) found that while one of these immunity genes had transcription levels that were significantly correlated across development, expression of the other gene was genetically decoupled. Although the authors interpreted the strong genetic correlation between larval and adult expression for one of the AMP genes as contradicting the ADH, they also acknowledged that they lacked a priori predictions regarding expected genetic correlations for these immune traits. One potential explanation for these contrasting decoupling patterns is that AMPs with decoupled expression are optimized for stage-specific pathogens, whereas coupled AMPs respond to pathogens that attack multiple life stages. Similarly, because some parasites and pathogens of *N. lecontei* are highly stage-specific (Coppel and Benjamin 1955; Wilson et al. 1992), we would expect at least some—but not necessarily all—immune genes to have decoupled gene expression. Consistent with these predictions, a manually curated set of *N. lecontei* immune-related genes (Vertacnik et al., in prep) had highly variable decoupling patterns across life stages (Supplemental Tables 3-5).

The final trait we consider that may frequently evolve under conflicting selection pressures across development is pigmentation. In insects, coloration is subject to diverse selection pressures—including abiotic factors (thermoregulation, UV resistance, desiccation tolerance), predation, and sexual selection—that are likely to change over the course of development (Medina et al., 2020; True, 2003). Consistent with the ADH, an analysis of larval and adult coloration in 246 butterfly species found that color evolution is strongly decoupled across butterfly development (Medina et al., 2020). Notably, whereas selection stemming from predation appears to constrain larval color evolution, sexual selection on adult males gives rise to extensive interspecific variation in adult color. Although Medina et al. (2020) did not investigate the genetic underpinnings of color trait decoupling, genetic analysis in another lepidopteran (*Bicyclus anynana*) demonstrates that melanism in larvae and adults is controlled by separate loci (Saenko et al., 2012). Similarly, our results suggest that pine sawflies have pronounced decoupling of gene-expression traits related to pigmentation across development. For example, among our top differentially expressed genes among different life stages were top candidate genes identified by a QTL mapping analysis of among-population variation in melanin-based *(pale)* and carotenoid-based *(Cameo2)* larval pigmentation traits (Linnen et al., 2018).

Together with previous studies, our findings are consistent with the ADH prediction that selection for optimal trait decoupling across development yields strikingly heterogeneous patterns of decoupling for ecologically important traits. Additional studies are needed, however, to demonstrate that highly and minimally decoupled traits evolve under stage-specific and stage-independent selection pressures, respectively. Nevertheless, this collection of studies highlights the importance of taking ecology and stage-specific selection pressures into account when testing predictions of the ADH.

### Comparing decoupling across development to decoupling between the sexes

Beyond metamorphosis, the logic of the ADH applies to any situation in which pervasive antagonistic selection favors the evolution of reduced genetic correlations. Although theories explaining sexual dimorphism are conceptually similar to the ADH, the evolutionary trajectories of sexually antagonistic selection may differ predictably from those under ontogenetically antagonistic selection. However, few studies have directly compared patterns of sex-biased and stage-biased gene expression. In one notable exception, Perry et al. (2014) evaluated how patterns of sex-biased expression in gonadal tissue change between three developmental stages of *Drosophila melanogaster*. In contrast to our finding that sex-biased expression was modest compared to stage-biased expression (Figure 2), they reported greater decoupling between the sexes than between larvae and prepupae. This apparent discrepancy could be explained by minimal antagonistic selection between dispersing *Drosophila* larvae and prepupae, the lack of a distinct molt between these observed stages, or tissue specificity. By contrast—and consistent with our findings—a comprehensive analysis of gene expression in different developmental stages, sexes, and castes of two ant species revealed that developmental stage had the largest impact on gene-expression profiles (Ometto et al., 2011). Given that the intensity of sexually antagonistic selection is likely to be highly variable across taxa, traits, and tissues (Connallon & Clark, 2013; Connallon, Débarre, & Li, 2018; Pennell & Morrow, 2013), studies in diverse taxa will be needed to evaluate if decoupling of stage-biased expression consistently exceeds sex-biased expression.

A final layer of complexity is that if the genetic targets and/or intensity of sexually antagonistic selection vary across an organism’s life cycle, patterns of sex-biased gene expression may also be decoupled over ontogeny. In support of this prediction, Perry et al. (2014) reported that 18-30% of sex-biased genes exhibited stage-specific sex-biased expression, with 4.5% of genes having opposing sex bias at different stages. Importantly, sex- and stage-specificity impacted evolutionary rates with male-biased genes evolving most rapidly when expressed throughout development while female-biased genes evolved most rapidly when expressed in larvae only. Overall, these results are consistent with the view that sexually antagonistic and stage-specific selection can lead to predictable differences in patterns of gene expression and molecular evolution.

### Limitations and future directions for ADH research

Although the patterns of gene-expression decoupling we observed in *N. lecontei* are consistent with the ADH, additional data are needed to: (1) verify that decoupled gene-expression phenotypes are independent at the genetic level, (2) demonstrate that decoupled gene-expression traits contribute to stage-specific adaptations, and (3) establish that metamorphosis is an adaptation for trait decoupling. Here, we discuss strategies for evaluating each of these additional ADH predictions.

First, if stage-specific levels of expression for a particular gene are genetically independent, alleles that contribute to expression variation in one life stage should not have pleiotropic effects on expression in another life stage (and vice versa). One way to evaluate this prediction is to perform quantitative trait locus (QTL) mapping on gene-expression traits at different life stages. Genetic independence would be supported if QTL for gene-expression traits measured at different stages are non-overlapping (e.g., (Freda et al., 2017; Saenko et al., 2012)). One example of this approach is a 2017 study that investigated genetic decoupling of thermal hardiness between larval and adult *D. melanogaster* (Freda et al., 2017). Whereas *D. melanogaster* larvae live in thermally stable rotting fruits and are only present in the warm months, flying adults experience a more variable thermal environment and are exposed to low temperatures during the overwintering generation. Consistent with these strong opposing selection pressures, thermal hardiness is completely decoupled across metamorphosis in this species. Moreover, decoupling of the thermal-hardiness phenotype is mirrored at a genetic level: loci that contribute to cold hardiness in larvae do not appear to have pleiotropic effects on adults, and vice versa (Freda et al., 2017). Interpreted in light of fruit-fly ecology, decoupled thermal hardiness phenotypes and alleles provide strong support for the ADH.

Second, the prediction that decoupled gene-expression traits contribute to stage-specific adaptation could be evaluated using multiple complementary approaches. For example, if decoupled genes contribute disproportionately to adaptation, genes exhibiting the most stage-biased expression patterns should also reveal a history of positive selection (e.g., evidence of recent selective sweeps or elevated rates of non-synonymous substitutions relative to the rest of the genome) (Vitti, Grossman, & Sabeti, 2013). Although this prediction has been confirmed by several studies for sex-biased genes (Assis et al., 2012; Drosophila 12 Genomes et al., 2007; Mank, Nam, Brunstrom, & Ellegren, 2010; Proschel et al., 2006; L. Yang, Zhang, & He, 2016), it has rarely been tested in the context of stage-biased expression across metamorphic boundaries (but see (Perry et al., 2014)). An alternative approach would be to use experimental genomics to connect genetic variants directly to fitness at different life stages (e.g., (Egan et al., 2015; Gloss, Groen, & Whiteman, 2016; Gompert et al., 2019; Ingvarsson, Hu, Lei, & de Meaux, 2017)). Following exposure to a selection regime that favors different traits at different ontogenetic stages, the ADH predicts that genes with the most decoupled expression will exhibit the most pronounced allele frequency shifts.

Third, to more directly test the hypothesis that metamorphosis itself is an adaptation for optimizing trait decoupling, comparative data can be used two evaluate two additional predictions: (1) metamorphosis is favored under ecological conditions that result in pervasive antagonistic pleiotropy across the life cycle and (2) metamorphosis facilitates trait decoupling. To disentangle the ecological and genetic correlates of metamorphosis from shared phylogenetic history, comparative tests of the ADH should focus on lineages that contain multiple independent origins of particular metamorphic phenotypes. For example, within holometabolous insects, hypermetamorphosis has evolved multiple times (Belles, 2011). Likewise, gains and losses of complex life cycles have been demonstrated in numerous taxa and are particularly well documented in insects and amphibians (Badets & Verneau, 2009; Bonett & Blair, 2017; Emmanuelle, Gwenaelle, & Armelle, 2010; Moran, 1994; Poulin & Cribb, 2002; Wiens, Kuczynski, Duellman, & Reeder, 2007).

## Conclusions

Overall, our transcriptomic analysis of a hypermetamorphic and sexually dimorphic sawfly demonstrates that patterns of gene-expression decoupling can be surprisingly predictable. These findings shed light on seemingly contradictory results reported in previous tests of the ADH and set the stage for follow-up studies on the genetic basis of stage-specific adaptation. However, rigorously testing the ADH and better understanding its relation to sexual dimorphism will ultimately require analyses of gene-expression decoupling in diverse taxa that vary in metamorphic and sexually dimorphic phenotypes. To gain maximal insight from decoupling analyses in other taxa, *a priori* predictions derived from in-depth knowledge of organismal ecology are essential. Although much work remains, these data are critical to understanding why metamorphosis is one of the most successful developmental strategies on the planet.

## Acknowledgements

For assistance with collecting and maintaining sawfly colonies, we thank current and past members of the Linnen laboratory. For constructive comments on versions of this manuscript, we thank Emily Bendall and Kathryn Everson. For computing resources, we thank the University of Kentucky Center for Computational Sciences and the Lipscomb High Performance Computing Cluster. Funding for this research was provided by the Kentucky Science and Education Foundation (KSEF-3492-RDE-019; to C.R.L.), the United States Department of Agriculture National Institute of Food and Agriculture (2016-67014-2475; to C.R.L.), and the National Science Foundation (DEB-CAREER-1750946; to C.R.L.).

## Data Accessibility Statement

All raw reads will be added to NCBI short read archive (accession numbers will become available upon peer-review publication). Gene expression data will be available through the Dryad repository (available upon peer-review publication).

## Author Contributions

DKH, KLV, and CRL participated in the study design. DKH, KLV, and ARK participated in the data collection. DKH and CRL analyzed the data. DKH and CRL drafted the manuscript. All authors read and approved the final manuscript.

## Notes

### Competing Interest Statement

The authors have declared no competing interest.

### Summary of Updates

After feedback, we have updated the manuscript.

